# A TAK1 Cytokine Toxicity Checkpoint Controls Anti-Cancer Immunity

**DOI:** 10.1101/2025.05.09.652721

**Authors:** Tirta M Djajawi, Anne Huber, Sarahi Mendoza Rivera, Akash Srivaths, Maede Salehi, Gokhan Gunay, Chloe Gerak, Liam Neil, Oliver Ozaydin, Olivia Voulgaris, Aleen Al Halawani, Dáire Gannon, Nooshin Khoshdoozmasouleh, Laura J Jenkins, Kok Fei Chan, Andreas Behren, Matthias Ernst, Lisa A Mielke, Emily J Lelliott, Hao Dong, Rebecca Feltham, Vivien R Sutton, Joseph A Trapani, John M Mariadason, Bhupinder Pal, Seamus J Martin, Stephin J Vervoort, Conor J Kearney

## Abstract

The success of cancer immunotherapies is currently limited to a subset of patients, which underscores the urgent need to identify the processes by which tumours evade immunity. Through screening a kinome-wide CRISPR/Cas9 sgRNA library we identified *MAP3K7* (TAK1) as a suppressor of CD8+ T cell mediated killing. We demonstrate that TAK1 acts as a cancer-intrinsic checkpoint by integrating signals from T cell-secreted TNF and IFNy effector cytokines to elicit a cytoprotective response. This cytoprotective response profoundly limits the anti-cancer activity these key effector molecules and completely abrogates bystander killing by perforin deficient T cells. Inhibition of the TAK1 checkpoint effectively redirects the combined TNF/IFNy pathway activation to promote inflammatory cell death via RIPK1 and Caspase-8 and simultaneously amplifies the output of the IFNy pathway, thereby priming cells for cytokine-induced cell death. Mechanistically, TAK1 deficiency led to proteasomal degradation of cFLIP, enhancing the formation of Complex II and subversion of other cytoprotective responses. Targeting the TAK1 checkpoint led to profound attenuation of tumour growth in immune competent mice, with minimal impact in immune deficient counterparts. Adoptive cell therapy led to preferential elimination of TAK1 deficient clones. Collectively, our study uncovers a cancer-intrinsic checkpoint controlled by TAK1 activity that switches TNF and IFNy responses from cytoprotective to apoptosis. Cancer cells exploit this to limit cell death in the presence of the cytotoxic lymphoctye cytokines TNF and IFNγ and therapeutic intervention can fully unleash the impact of these effector molecules both on the direct target and bystander cells. These findings highlight the clinical development of TAK1 biologics as a potential strategy to improve cancer immunotherapies through harnessing and enhancing the cytotoxic potential of CTL-derived cytokines.

**In Brief:** Djajawi et al. identify TAK1 as a cancer-intrinsic cytokine toxicity checkpoint that limits the efficacy of anti-cancer immune responses. Cancer cells exploit TAK1 activity to confers resistance to CD8+ T cell-derived TNF and IFNγ-induced apoptosis, limiting both direct and bystander killing. Mechanistically, TAK1 loss acts through 1. cFLIP and RIPK1 to promote cell death, 2. The inactivation of pro-survival signals and 3. Via amplification of the IFNy response. Targeting of this TAK1 checkpoint enhanced anti-tumour immunity in vivo and improved adoptive cellular therapy. These findings identify a strategy employed by transformed cells to avoid destruction by inflammatory cytokines and provide new therapeutic vulnerabilities for enhancing immunotherapies.

**Highlights:**

- **CRISPR screens identify TAK1 as a tumour-intrinsic survival checkpoint limiting destruction by CD8+ T cells.**
- **TAK1 protects tumour cells from combined TNF and IFNγ-induced apoptosis.**
- **TAK1 loss destabilizes cFLIP, priming cells for STAT1 and RIPK1-dependent cell death.**
- **TAK1 loss promotes tumour control in immune competent animals and enhances the efficacy of adoptive cell therapy.**

## Introduction

Immune checkpoint blockade (ICB) has reshaped the clinical management of multiple cancers, particularly melanoma, non-small cell lung cancer, and renal cell carcinoma, by unleashing the cytotoxic potential of T cells against tumour cells. Antibodies targeting programmed cell death protein 1 (PD-1), its ligand PD-L1, or cytotoxic T-lymphocyte-associated protein 4 (CTLA-4) restore anti-tumour T cell function and can induce durable clinical responses in a subset of patients (1, 2). However, overall response rates remain limited, with the majority of patients exhibiting primary or acquired resistance (3, 4). These clinical limitations underscore the pressing need to dissect resistance to either tumour-intrinsic mechanisms or checkpoints that govern resistance to T cell–mediated killing (5).

Effective anti-tumour immunity requires not only the ability of CD8+ T cells to recognize tumour antigens presented via major histocompatibility complex class I (MHC-I), but also the execution of cytotoxic functions through cytokine secretion and direct induction of cell death (6). Two central T cell–derived cytokines, tumour necrosis factor (TNF) and interferon gamma (IFNγ), play pivotal roles in shaping tumour cell fate (7). IFNγ enhances antigen presentation, activates pro-inflammatory transcriptional programs, and modulates susceptibility to apoptosis (8, 9), while TNF signals through TNFR1 to induce either pro-survival NF-κB signaling or cell death, depending on the cellular context (10, 11). Resistance to either cytokine is becoming increasingly realised as a key driver immunotherapy failure (12, 13).

Large-scale CRISPR-based functional screens have emerged as powerful tools for identifying genes that govern tumour–immune interactions. Pioneering studies have systematically interrogated tumour cell vulnerabilities in the context of cytotoxic T cell pressure and defined core immune resistance mechanisms involving antigen presentation, IFNγ signaling, and TNF regulation (14–16). Collectively, these studies have demonstrated that CRISPR screening in antigen-expressing tumour cells under T cell selection pressure can reveal positive and negative regulators of antigenicity and immune recognition, highlighting the value of focused *in vitro* or *in vivo* immune co-culture systems for decoding tumour-intrinsic immune evasion. However, the statistical dominance of well-characterized pathways in whole-genome datasets may obscure additional mechanisms of resistance that could represent untapped therapeutic targets (17).

To identify genes that protect tumour cells from CTL attack, we custom cloned a focused sgRNA library targeting the kinome (2 sgRNAs per gene). We then performed a CRISPR/Cas9 screen whereby B16F10 cells carrying the library were exposed to the rounds of killing by antigen matched OT-I CD8+ T cells. Here we identified Transforming Growth Factor Beta-Activated Kinase 1 (TAK1, MAP3K7) as a key gene that protects B16F10 cells from CTL attack. TAK1 is a serine/threonine kinase that functions downstream of TNFR1, Toll-like receptors (TLRs), and IL-1R, integrating stress and inflammatory signals through activation of NF-κB and MAPK cascades (18, 19). Although TAK1 is well-established as a mediator of immune and developmental signaling, its role in regulating tumour sensitivity to immune effector cytokines has remained undefined.

We found that loss of TAK1 sensitizes tumour cells to cell death in response to combined TNF and IFNγ exposure, released by cytotoxic lymphocytes. Mechanistically, TAK1-deficient cells exhibit proteasome-dependent degradation of the caspase-8 inhibitor cFLIP following TNF stimulation, removing a key checkpoint that restrains extrinsic apoptosis. This degradation event lowers the threshold for IFNγ-induced killing, which is further amplified by STAT1-dependent transcriptional programs.

To define the molecular effectors of this cytokine-induced death, we conducted genome-wide CRISPR counter-screens in a TAK1 knockout background, revealing that RIPK1 and STAT1 are essential mediators of apoptosis under these conditions. These data support a model in which RIPK1 and STAT1 function cooperatively to drive cell death when TAK1 is absent or inhibited.

Collectively, these observations define a multi-layered tumour-intrinsic checkpoint centred around TAK1, cFLIP, RIPK1 and STAT1 that integrates inflammatory signals and intracellular stress responses to suppress cell death in the context of T cell–mediated attack. Here we uncover novel regulators of immune resistance and identify new candidate targets for combination immunotherapy approaches aimed at disrupting tumour cell survival checkpoints under inflammatory stress driven by CTL attack.

## Results

### Kinome-wide CRISPR screens identify TAK1 as a suppressor of CD8+ T cell-mediated killing

To systematically identify kinases that mediate tumor-intrinsic resistance to cytotoxic T lymphocyte (CTL)-mediated immune pressure, we performed a focused CRISPR-Cas9 screen using a custom sgRNA library targeting all annotated kinases. B16F10-OVA melanoma cells stably expressing Cas9 were transduced with the pooled lentiviral kinase library and subjected to three consecutive rounds of overnight co-culture with activated, antigen-specific OT-I CD8+ T cells (Figure 1A). Surviving tumor cells were collected after each round and analyzed by next-generation sequencing to quantify sgRNA representation relative to baseline untreated populations. As can be seen from Figure 1B, subsequent rounds of OT-I exposure led to a loss of sgRNA library complexity, as evident from an increasing Gini-Index (20).

**Figure 1.**
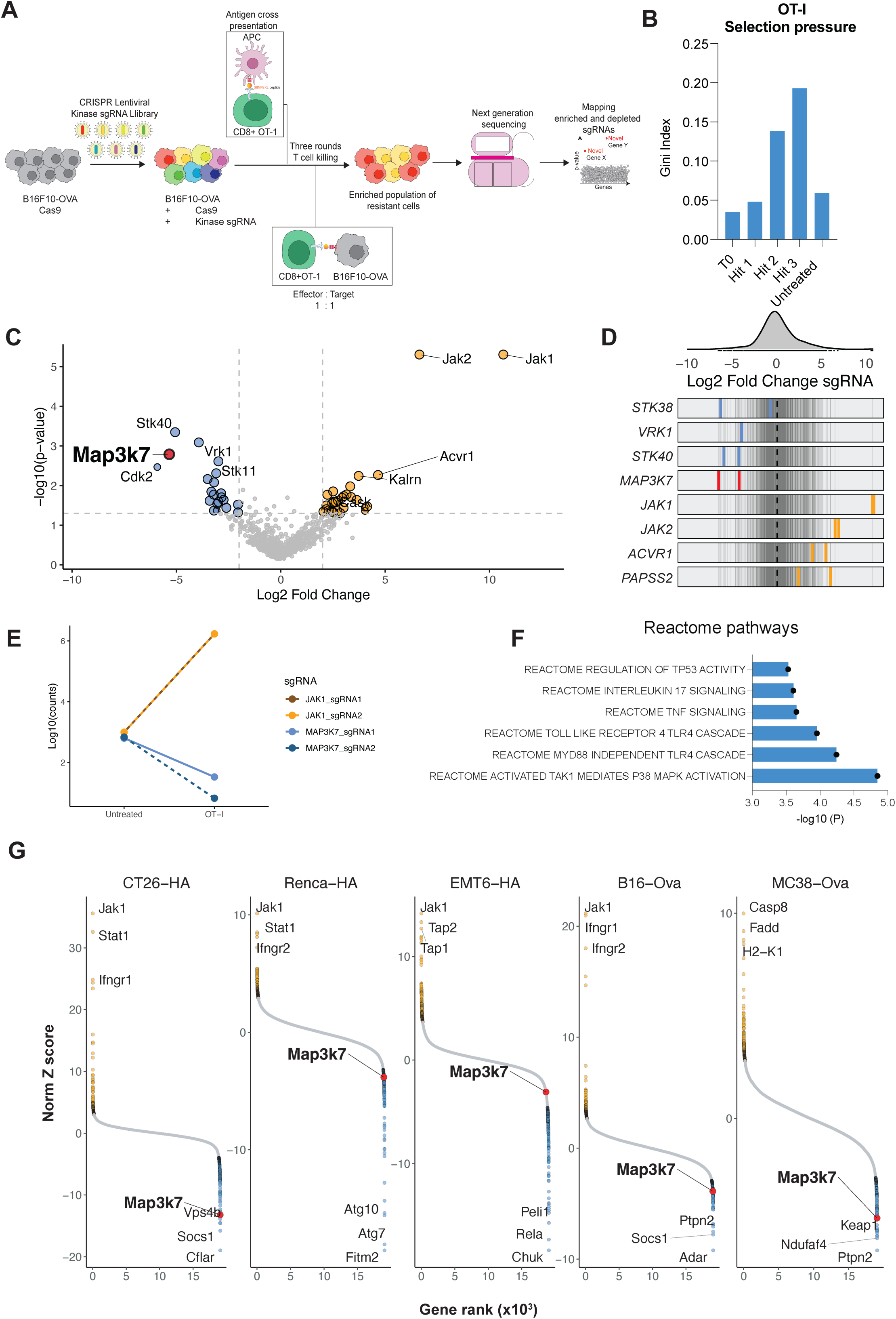
Kinome-wide CRISPR Cas9 screen reveals that loss of TAK1 promotes CD8+ T cell killing. (A) Schematic of the experimental workflow. B16F10-OVA cells expressing Cas9 were transduced with a lentiviral sgRNA library targeting the mouse kinome and subjected to three rounds of co-culture with OT-I CD8+ T cells (E:T ratio 1:1). Surviving tumor cells were harvested and sgRNA abundance quantified by next-generation sequencing. (B) The Gini-index is displayed throughout three rounds of OT-I T cell exposure. (C) Volcano plot showing enriched and depleted sgRNAs from screen depicted in Figure 1A. The p-values were generated by MaGeCK analysis. (D) Distribution of selected sgRNA in response to OT-I selection pressure, highlighting both enriched and depleted sgRNAs relative to the untreated population. (E) Enrichment/depletion of sgRNAs targeting JAK1 and MAP3K7 in the untreated condition and OT-I treated condition. (F) Summary of gene ontology for the most affected pathway from kinome-wide CRISPR Cas9 screen. Reactome pathways affecting TAK1 were significantly altered. (G) Gene-level CRISPR screen results across six tumor models (CT26-HA, Renca-HA, EMT6-HA, B16F10-OVA, and MC38-OVA), showing log₂ fold-change in sgRNA abundance following T cell co-culture from Lawson et al, (12). The MAP3K7 is highlighted as a red dot.

Analysis of sgRNA enrichment and depletion revealed distinct patterns of selection, with multiple genes showing significant changes in representation following T cell selection pressure (Figure 1C-E). Notably, sgRNAs targeting MAP3K7 (TAK1) were among the most significantly depleted, suggesting these kinases are critical for tumor cell survival in the context of immune attack. As expected, sgRNAs sgRNAs targeting JAK1 and JAK2 were heavily enriched, confirming optimal screen performance (Figure 1C-E). Indeed, Reactome pathway analysis revealed depletion of genes in pathways related to P38-mediated TAK1 activation as well as TNF signalling (Figure 1F)

To assess the generalizability of these findings, we analysed similar published screening data across multiple tumor models expressing defined antigens, including CT26-HA, Renca-HA, EMT6-HA, MC38-OVA, and B16F10-OVA (12). *MAP3K7* was consistently depleted across all tumor types, reinforcing its conserved role in protecting tumor cells from T cell-mediated cytotoxicity (Figure 1G). Similarly, enrichment of JAK1 was observed across multiple cell lines, highlighting the broader relevance of JAK-STAT signaling in tumour immune evasion. These data argue that TAK1 plays a key role in tumour-intrinsic defence against T cell attack and establish a robust platform for identifying therapeutic vulnerabilities under immune selection.

### TAK1 protects tumours from CTL-derived TNF and IFNy

To validate TAK1 as a key resistance gene against CTL attack, we generated TAK1-deficient B16F10-OVA melanoma cells using three independent sgRNAs targeting MAP3K7. Western blotting confirmed efficient depletion of TAK1 protein across all sgRNAs (Figure 2B). To determine whether TAK1 loss altered tumor cell susceptibility to CD8+ T cell–mediated cytotoxicity, we utilized a fluorescent competition assay wherein wild-type (WT) B16F10 cells expressing BFP were mixed at a 1:1 ratio with TAK1-knockout cells expressing mCherry. Mixed populations were co-cultured with antigen-specific OT-I CD8+ T cells for 48 hours (Figure 2A). Flow cytometry revealed a striking and reproducible reduction in the proportion of mCherry⁺ TAK1-deficient cells relative to BFP⁺ WT cells following T cell exposure, indicating selective elimination of TAK1-deficient tumor cells (Figure 2C-D). We observed similar results using CT26-OVA cells and MC38-OVA cells (Supplementary Figure 1A-B). Indeed, re-introduction of a FLAG-tagged TAK1 cDNA rescued the phenotype, confirming the specificity of the system (Figure 2E-F). To determine whether this phenotype was dependent on TAK1 kinase activity, we reconstituted TAK1-deficient cells with either wild-type or kinase-dead (D175A) Flag-tagged TAK1 constructs. Ectopic expression of wild-type TAK1 fully rescued the survival defect in the presence of OT-I T cells, whereas the D175A mutant failed to restore resistance (Figure 2G-H). These findings indicate that the kinase function of TAK1 is required to protect tumor cells from T cell-induced death.

**Figure 2.**
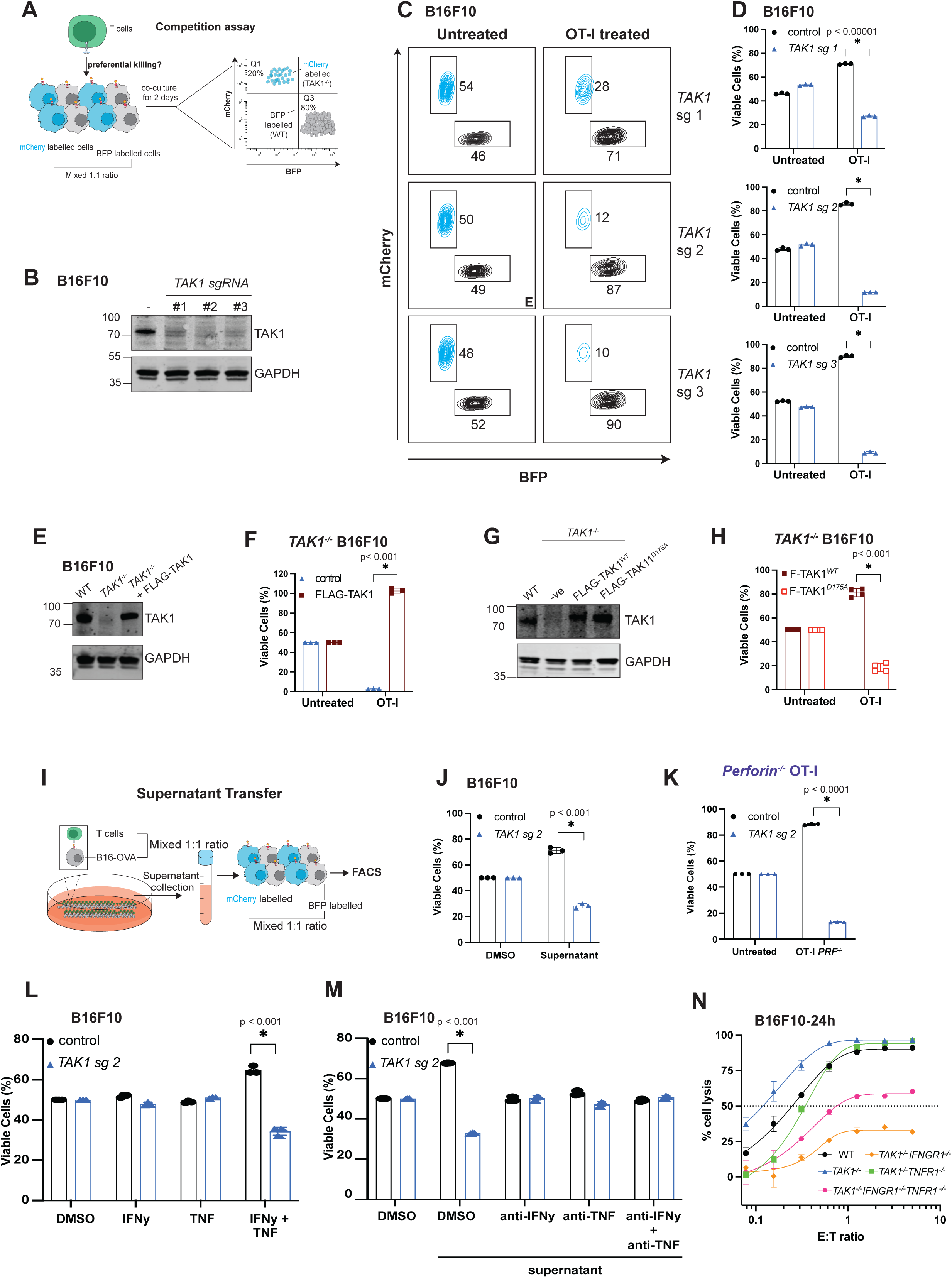
TAK1 deficient melanoma cells are sensitive to combined cytokine treatments of IFNy and TNF. (A) Schematic of the competition assay. B16F10-OVA cells labeled with mCherry (TAK1 knockout) and BFP (wild-type) were mixed at a 1:1 ratio and co-cultured with OT-I CD8+ T cells for 48 hours. Preferential elimination of TAK1-deficient cells was assessed by flow cytometry. (B) TAK1 was targeted with three independent sgRNAs in B16F10 cells using CRISPR RNPs, western blots are displayed to confirm knockout. (C-D) (C) Representative FACS plots of mixed BFP⁺ (WT) and mCherry⁺ (TAK1^⁻/⁻^) cells before and after co-culture with OT-I T cells for 48 hours, showing selective depletion of TAK1-deficient cells. (D) Bar chart derived from (C) showing that TAK1 deficient B16F10-OVA were preferentially killed by OT-I CD8+ T cells. Data were analysed by unpaired t-test for three replicates, *p values <0.00001. (E-F) (E) Western blot confirming TAK1 depletion by sgRNAs and re-expression of FLAG-TAK1. (F) The resulting competition assay is depicted from *TAK1^-/-^* B16F10-OVA cells or ones that express FLAG-TAK1 after no treatment or after OT-I T CD8+ cells for 48 hours. Data were analysed by unpaired t-test for three replicates, * p value <0.001. (G-H) (G) Western blot confirming TAK1 depletion by sgRNAs and re-expression of either FLAG WT TAK1 or D175A TAK1. The resulting competition assay is depicted in (H). Data were analysed by unpaired t-test for three replicates, * p value <0.001. (I) Schematic of supernatant transfer assay that is derived from OT-I CD8+ T cells and B16F10-OVA co-culture. (J) Supernatant transfer assay: conditioned media from OT-I/B16 co-cultures was sufficient to kill TAK1-deficient cells, an effect reversed by IFNγ/TNF neutralization or receptor loss. (K) Competition assay as described in Figure 2A, using Perforin deficient OT-I CD8+ T cells. Data were analysed by unpaired t-test for three replicates, * p value <0.0001. (L) Competition assay as described in Figure 2A, using cytokines treatment instead of OT-I CD8+ T cells. Wild type or TAK1 knockout B16F10 cells were exposed to IFNy (1 ng/mL), TNF (20ng/mL), or both cytokines for 48 hours. Data were analysed by unpaired t-test for three replicates, * p value <0.001. (M) Competition assay with supernatant as described in Figure 2I-J in the presence or absence of IFNy or TNF neutralising antibodies. Data were analysed by unpaired t-test for three replicates, * p value <0.001. (N) OT-I killing assay using B16F10 targets at the indicated effector to target ratios where the indicated genes (*TAK1, IFNGR1, TNFR1)* had been deleted using CRISPR. Cell killing was measured 24h after T cell exposure.

To test whether T cell-derived cytokines were sufficient to mediate this effect, we performed a series of supernatant transfer assays. Conditioned media from OT-I and B16F10-OVA co-cultures was collected and transferred onto untreated tumor cells. This supernatant induced substantial cell death in TAK1-deficient, but not WT, B16F10 cells (Figure 2I-J). Next, we assessed whether tumor cell killing required perforin-mediated cytolysis. Perforin knockout OT-I T cells retained their ability to preferentially kill TAK1-deficient B16F10 cells (Figure 2K), confirming that cytotoxicity occurred through a cytokine-driven mechanism. To directly test the effect of recombinant cytokines, TAK1-deficient and WT cells were treated with IFNγ and TNF, either alone or in combination. TAK1-deficient cells exhibited profound sensitivity to cytokine-induced killing, with maximal cell death observed under combined IFNγ and TNF treatment (Figure 2L). We observed similar results across a variety of cell lines using either genetic deletion of MAP3K7 or inhibition of TAK1 using Takinib (Supplementary Figure 1C). These data demonstrate that TAK1 is essential for tumor cell survival in the presence of inflammatory cytokine signaling. Neutralization of IFNγ and TNF with blocking antibodies significantly reduced this effect, confirming that soluble cytokines secreted by CD8⁺ T cells are both necessary and sufficient to drive tumor cell death in the absence of TAK1 signaling (Figure 2M).

Killing assays across a range of effector-to-target (E:T) ratios demonstrated significantly increased lysis of TAK1-deficient cells compared to WT controls (Figure 2N). Notably, combined deletion of the IFNγ receptor (IFNGR1) and the TNF receptor (TNFR1) (Supplementary Figure 1D) in TAK1-deficient cells largely abrogated T cell-mediated cytotoxicity, implicating inflammatory cytokines, rather than perforin or granzyme-mediated lysis, as the dominant drivers of tumor cell death in the absence of TAK1 (Supplementary Figure 1C). Together, these results delineate a non-canonical mode of tumor clearance by CD8⁺ T cells in which the inflammatory cytokines IFNγ and TNF, act in concert to eliminate tumor cells lacking TAK1 signaling.

### TAK1 is required for NF-κB–driven transcriptional programs but dispensable for IFNγ-mediated gene induction

To investigate how TAK1 controls cytokine-driven gene expression, we conducted RNA-sequencing on wild-type and TAK1-deficient B16F10 cells following stimulation with TNF, IFNγ, or their combination. In WT cells, TNF, IFNγ, and combined TNF+IFNγ treatment induced substantial transcriptional reprogramming, with the strongest response observed in the combination treatment (Figure 3A-D).

**Figure 3.**
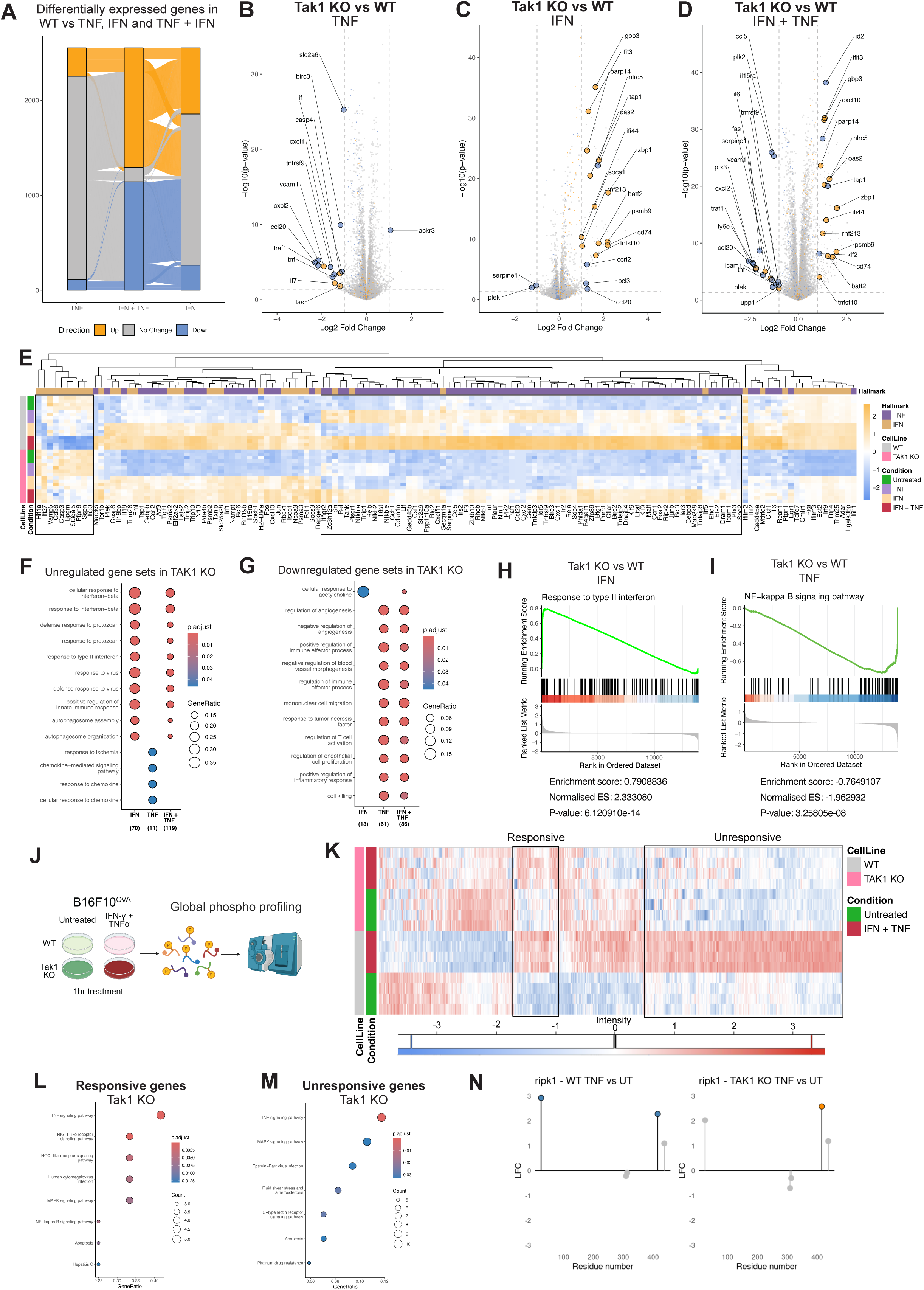
(A) Number of differentially expressed genes in WT B16F10 cells following treatment with TNF, IFNγ, or the combination (TNF + IFNγ). Genes are grouped by directionality (upregulated, downregulated, no change). (B-D) Volcano plots showing differential gene expression between TAK1 KO and WT B16F10 cells following IFNγ (B), TNF (C), or TNF + IFNγ (D) treatment. Key inflammatory and immune-related genes are labelled. (E) Heatmap of cytokine-induced gene expression across four conditions (untreated, TNF, IFNγ, TNF+IFNγ) in WT and TAK1 KO B16F10 cells. (F-G) Gene ontology (GO) enrichment analysis of genes upregulated *F*, or downregulated, *G*, in WT B16F10 cells treated with IFNγ, TNF, or both cytokines. (H-I) Gene Set Enrichment Analysis (GSEA) showing TAK1-dependent changes in hallmark pathways. (H) “NF-κB signaling” is significantly enriched in WT cells treated with TNF. (I) “Response to type II interferon” is enriched in KO cells treated with IFNγ. (J) Schematic overview of phosphoproteomics experiment. (K) Heatmap of global phosphoproteomic profiling of WT and TAK1 KO B16F10 cells treated with TNF+IFNγ for 1 hour. Phosphosite intensity is shown for each condition. (L-M) KEGG pathway enrichment for phosphosites that are TAK1 dependant and TAK1 independent. (N) Phophosite intensity changes in RIPK1 comparing wild type and TAK1 knockout cells as described in *A*.

To determine the contribution of TAK1 to these responses, we compared transcriptomic profiles between WT and TAK1 KO cells. TNF-induced gene expression was markedly impaired in TAK1 KO cells, with significant downregulation of canonical NF-κB target genes including Nfkbia, Relb, Ccl20, and Tnfaip3 (Figure 3E). In contrast, the transcriptional response to IFNγ was largely preserved in the absence of TAK1 (Figure 3E), suggesting TAK1 is not required for IFNγ-mediated STAT1 activation and gene induction. Consistent with this, combined TNF+IFNγ treatment in TAK1-deficient cells resulted in selective loss of NF-κB target gene expression, while IFN-stimulated genes such as Ifit3, Gbp3, and Irf1 remained upregulated (Figure 3E). Indeed, Gene ontology (GO) enrichment analysis revealed that upregulated pathways in TAK1 knockout cells included response to interferon while the downregulated pathways included response to TNF (Figure 3F-G). Furthermore, Gene set enrichment analysis (GSEA) further confirmed that the NF-κB signaling pathway was significantly suppressed in TAK1 KO cells following TNF treatment (Figure 3H). Conversely, genes involved in the response to type II interferon were enriched in TAK1-deficient cells (Figure 3I).

### TAK1 selectively regulates phosphorylation programs downstream of TNF and IFN**γ.**

Next, we performed global phosphoproteomic profiling of Wild type and TAK1 KO cells treated with IFNγ+TNF for 1 hour (Figure 3 J). Unsupervised clustering revealed distinct phospho-signatures in TAK1-deficient cells, and gene ontology analysis of regulated phosphoproteins identified enrichment for immune signaling, cytokine response, and cell death pathways (Figure 3H–K). In particular, phosphosites involved in NF-κB and MAPK signaling—such as those on Rela, Ripk2, and Map3k8— were reduced in TAK1 KO cells, while STAT1-associated phosphorylation events remained intact (Figure 3K). We stratified phosphosites as TAK1-sensitive or TAK1-independent based on their responsiveness to TNF+IFNγ in WT but not KO cells. KEGG enrichment of TAK1-sensitive sites highlighted pathways such as TNF signaling, RIG-I–like receptor signaling, and NF-κB activation (Figure 3L). In contrast, phosphosites maintained in TAK1 KO cells were enriched for MAPK and viral infection-related processes (Figure 3M). Indeed, we next examined site-specific phosphorylation changes on RIPK1 following TNF and IFNγ stimulation. Phosphorylation was induced in wild-type cells, but this response was selectively altered in TAK1-deficient cells. These data indicate that TAK1 is required for specific RIPK1 phosphorylation events, while others occur independently of TAK1. Together, this highlights a role for TAK1 in modulating RIPK1 signaling downstream of TNF and IFNy.

### A CRISPR resistance screen identifies genes mediating cell death in response to IFNγ and TNF in the absence of TAK1

Next we repeated the CRISPR screen using the kinome library using recombinant cytokines instead of CTL. As can be seen from Figure 4A, TAK1 was not among the top depleted hits upon treatment with TNF or IFNy alone. However, TAK1 was significantly depleted in the screen in which cells were treated with both cytokines.

**Figure 4.**
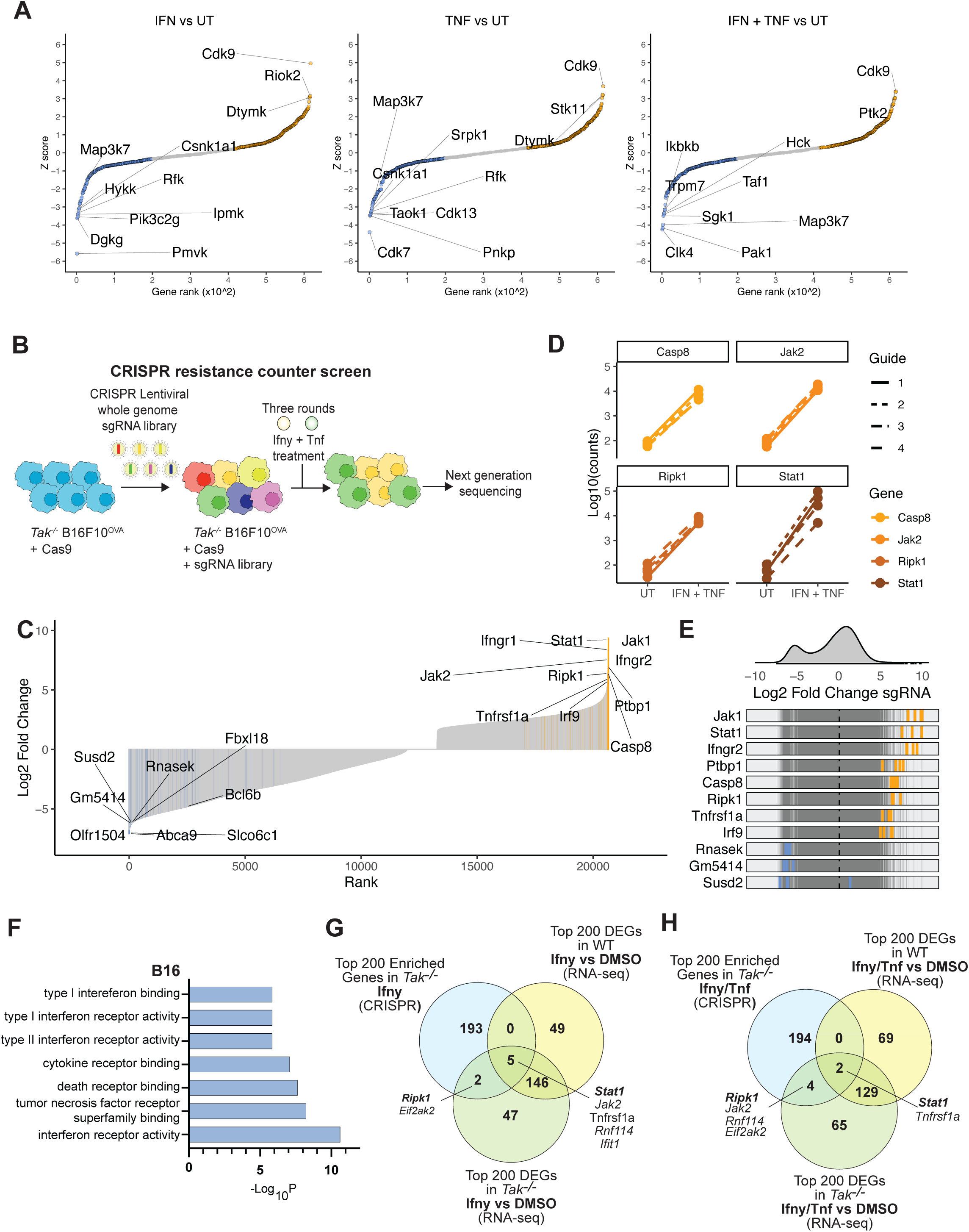
Deficiency in either interferon gamma or tumour necrosis factor signaling pathway can reverse TAK1 loss induced killing. (A) Wild type B16F10 cells expressing Cas9 used for a CRISPR screen using the kinome-wide library. Cells were left untreated or treated with TNF (20ng/ml), IFNy (1ng/ml) or both for 7 days. (B) Schematic of genome-wide CRISPR resistance screen in TAK1-deficient B16F10-OVA cells expressing Cas9. Cells were transduced with a lentiviral sgRNA library, treated with IFNγ (1ng/mL) and TNF (20 ng/mL) and subjected to next-generation sequencing to identify enriched sgRNAs. (C) Volcano plot showing enriched and depleted sgRNAs in *TAK1^-/-^* cells treated with IFNy + TNF, highlighting genes whose loss confers resistance or sensitisation. The p-values were generated by MaGeCK analysis. (D) Examples of enriched genes (CASP8, JAK2, RIPK1, STAT1) from Figure 4B. (E) Distribution of selected sgRNAs from Figure 4B, highlighting both enriched and depleted sgRNAs relative to untreated control. (F) Gene ontology (GO) term enrichment of CRISPR screen hits under IFNy + TNF, showing enrichment for cytokine receptor activity, death receptor binding, and interferon signaling pathways. (G-H) Venn diagrams comparing the top 200 CRISPR-enriched genes (Figure 4B) with top 200 differentially expressed genes (DEGs) identified by RNA-seq in WT or TAK1 KO cells treated with IFNγ (F) or IFNγ +. TNF (G).

To uncover genetic dependencies that drive cell death in response to inflammatory cytokines in TAK1 null cells, we conducted a whole-genome CRISPR resistance screen in TAK1-deficient B16F10-OVA cells expressing Cas9 (Figure 4B). Cells were transduced with a lentiviral sgRNA library, treated with IFNγ and TNF, then subjected to next-generation sequencing to quantify sgRNA abundance post-selection (Figure 4B).

Enrichment analysis identified key resistance genes including RIPK1, CASP8 and STAT1 as top hits, highlighting these genes as key mediators of cell death in the absence of TAK1 upon cytokine exposure (Figure 4C-E). Interestingly, the endosomal trafficking regulator RNASEK further sensitised to cell death, suggesting that the autophagy machinery may co-operate with TAK1 to protect cells from TNF and IFNy (Figure 4C-E). Gene Ontology enrichment analysis of screen hits revealed significant overrepresentation of terms related to interferon receptor activity, death receptor binding, and cytokine receptor signaling (Fig. 4F), consistent with the central involvement of immune signaling pathways in mediating cell death in TAK1-deficient cells.

We next compared the top 200 sgRNA-enriched genes from the CRISPR screen with RNA-seq–defined cytokine-responsive transcriptional programs. A limited overlap was observed between enriched hits and differentially expressed genes (DEGs) in either WT or TAK1-deficient cells, suggesting that resistance to cell death is not solely driven by transcriptional induction (Figure 4G–H). Genes such as RIPK1, JAK2, STAT1, and RNF114 appeared in both datasets, implying dual roles in cytokine response and cell survival regulation.

Taken together, these data demonstrate that in the absence of TAK1, IFNγ and TNF elicit a cell death program dependent on RIPK1, STAT1, and CASP8, and reveal a cell death checkpoint controlled by interferon and TNF receptor-proximal signaling.

### TAK1 suppresses STAT1- and RIPK1-dependent inflammatory cell death downstream of IFNγ and TNF

To confirm that the cell death was caspase-mediated, we treated TAK1-deficient cells with the pan-caspase inhibitor Q-VD-OPh during cytokine stimulation. Q-VD entirely blocked IFNγ+TNF-induced cell death (Figure 5A), indicating a caspase-dependent mechanism of cell death.

**Figure 5.**
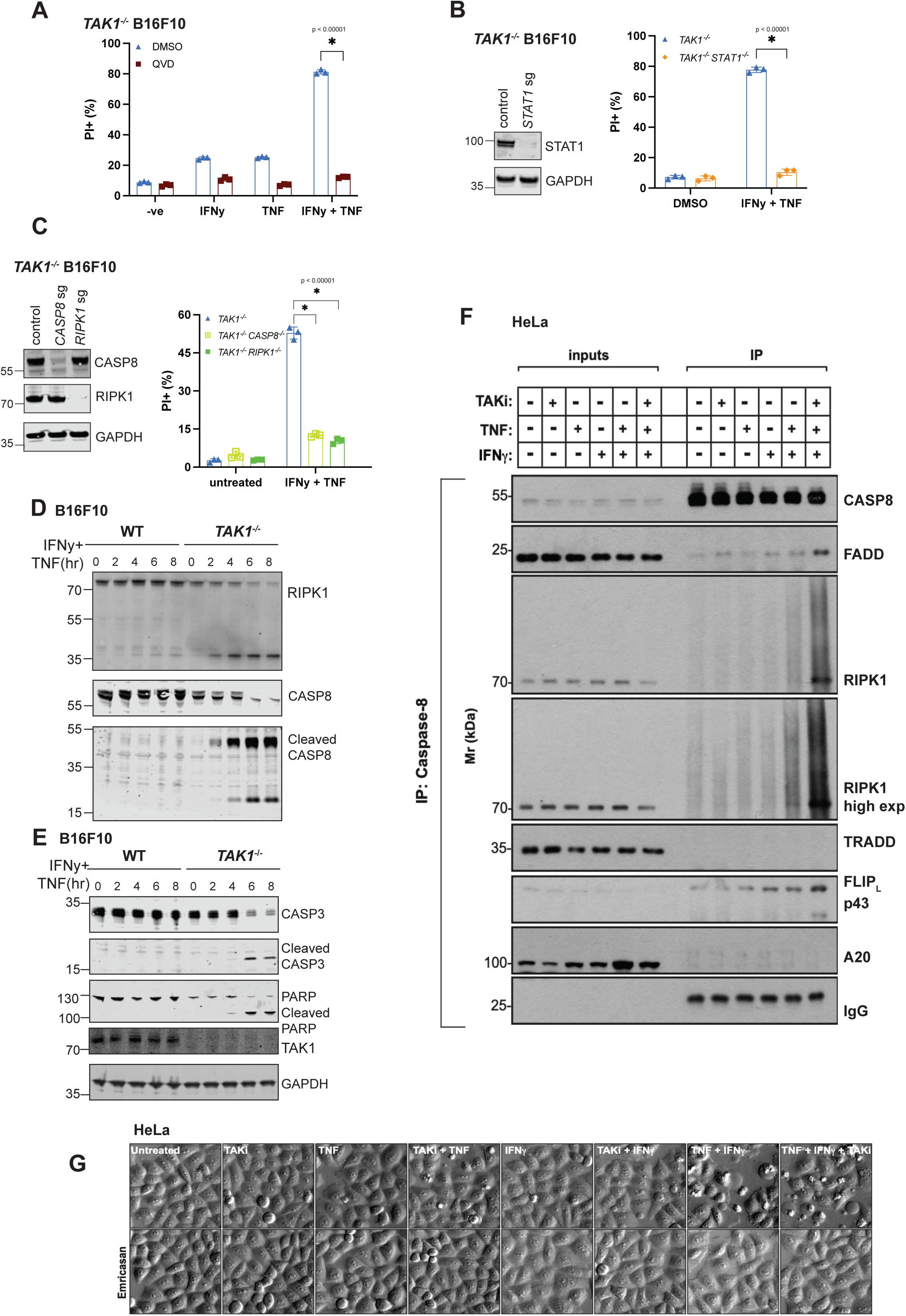
TAK1 limits inflammatory death mediated by STAT1 and RIPK1 in response to IFNy and TNF treatment. (A) IFNγ (1 ng/mL) and/or TNF (20 ng/mL)-induced cell death in *TAK1^⁻/⁻^* cells, with or without the pan-caspase inhibitor Q-VD-OPh (40µM). Data were analysed by unpaired t-test for three replicates, *p value < 0.00001. (B) Cell death assay (PI staining) of *TAK1^⁻/⁻^* and *TAK1^⁻/⁻^STAT1^⁻/⁻^* B16F10 cells following IFNγ and TNF treatment for 24 hours. Data were analysed by unpaired t-test for three replicates, *p value < 0.00001. (C) Propidium iodide (PI) staining of *TAK1^⁻/⁻^*, *TAK1^⁻/⁻^CASP8^⁻/⁻^*, and *TAK1^⁻/⁻^RIPK1^⁻/⁻^* B16F10 cells treated with IFNγ and TNF treatment for 24 hours. Data were analysed by unpaired t-test for three replicates, *p value < 0.00001. (D-E) Immunoblots of apoptotic markers over a time course of IFNγ and TNF treatment (0–8 hours) in WT and *TAK1^⁻/-^* B16F10*^⁻^*cells. TAK1-deficient cells show early and enhanced cleavage of caspase-8, caspase-3, and PARP, indicating caspase-mediated apoptosis. (F) Whole-cell lysates were prepared from HeLa cells treated with IFNγ (100 ng/mL), TNF (5 ng/mL), the TAK1 inhibitor (TAKi, 5 µM), or combinations as indicated. Caspase-8 was immunoprecipitated from cell lysates and analyzed by immunoblotting for the indicated binding partners. Inputs (left panels) show total protein levels of Caspase-8, FADD, RIPK1, TRADD, FLIPL, A20, and IgG (light chain control). Immunoprecipitates (IP: Caspase-8) (right panels) reveal inducible interactions between Caspase-8 and FADD, RIPK1, and TRADD upon cytokine stimulation in the presence of TAK1 inhibition. RIPK1 recruitment to the complex is strongly enhanced by combined IFNγ+TNF treatment with TAK1i. FLIPL and A20 are also modestly enriched under these conditions. High-exposure RIPK1 blot confirms increased interaction upon co-treatment. (G) Differential Interference Contrast microscopy for HeLa cells treated with IFNγ (100 ng/mL), TNF (2 ng/mL), the TAK1 inhibitor (TAKi, 5 µM), Emricasan (5 µM) or combinations as indicated. Images are representative of 3 independent experiments.

To determine whether the transcription factor STAT1 is required for cytokine-induced death in TAK1-deficient tumor cells, we generated *STAT1* knockout B16F10 cells in a TAK1-null background using CRISPR-Cas9 (Figure 5B). Indeed, *STAT1* deletion rescued the viability of TAK1-deficient cells following IFNγ+TNF treatment (Figure 5B). Next, we assessed whether key components of the extrinsic apoptosis pathway were involved in TNF+IFNy-induced cell death. Indeed, knockout of *CASP8* or *RIPK1* in TAK1-deficient B16F10 cells abrogated IFNγ+TNF-induced apoptosis, suggesting enhanced formation of the apoptosome under these conditions (Figure 5C). Furthermore, we observed enhanced cleavage of RIPK1, Caspase-8, Caspase-3 and PARP in the absence of TAK1 (Figure 5 D-E).

To investigate the molecular composition of the cell death-inducing complex that forms in the absence of TAK1 activity, we immunoprecipitated endogenous Caspase-8 from cells treated with IFNγ, TNF, and/or Takinib (TAKi). Immunoblotting revealed that under baseline conditions or single cytokine treatment, Caspase-8 associated weakly with FADD and minimally with RIPK1 or TRADD. However, co-treatment with IFNγ and TNF in the presence of TAK1 inhibition robustly promoted the assembly of a RIPK1–FADD–Caspase-8 complex (Figure 5F). RIPK1 co-immunoprecipitation was particularly strong under these conditions, consistent with its proposed role as a scaffold that facilitates Caspase-8 activation. TRADD was also recruited, suggesting upstream TNFR1 complex involvement. While the anti-apoptotic regulator FLIP_L_ was modestly present in all conditions, its levels did not increase substantially upon co-treatment. A20, a negative regulator of RIPK1 signaling, was weakly associated with the complex. These results indicate that TAK1 activity restrains the formation of a cytotoxic death-inducing complex centered on Caspase-8 and RIPK1, which becomes strongly assembled in the presence of inflammatory cytokines when TAK1 is absent or inhibited. We also confirmed enhanced apoptosis under these conditions using microscopy (Figure 5G).

### cFLIP limits caspase-8 activation and cell death in TAK1-deficient cells under inflammatory cytokine exposure

Next we hypothesised that cFLIP may be responsible for the increased sensitivity to TNF and Interferon gamma in the absence of TAK1. Again, we re-analysed data from Lawson et al and found that *CFLAR*, the gene encoding cFLIP was significantly depleted across all cell lines tested (Figure 6A). In wild-type B16F10 cells, cFLIP levels remained relatively stable upon stimulation with IFNγ and TNF. In contrast, TAK1-deficient cells exhibited a marked reduction in cFLIP_L_ protein levels following cytokine treatment, with levels progressively decreasing over time (Figure 6B). The most profound downregulation occurred in response to TNF alone (Figure 6C), suggesting that TNF-induced cFLIP degradation contributes to enhanced caspase-8 activity in the absence of TAK1.

**Figure 6.**
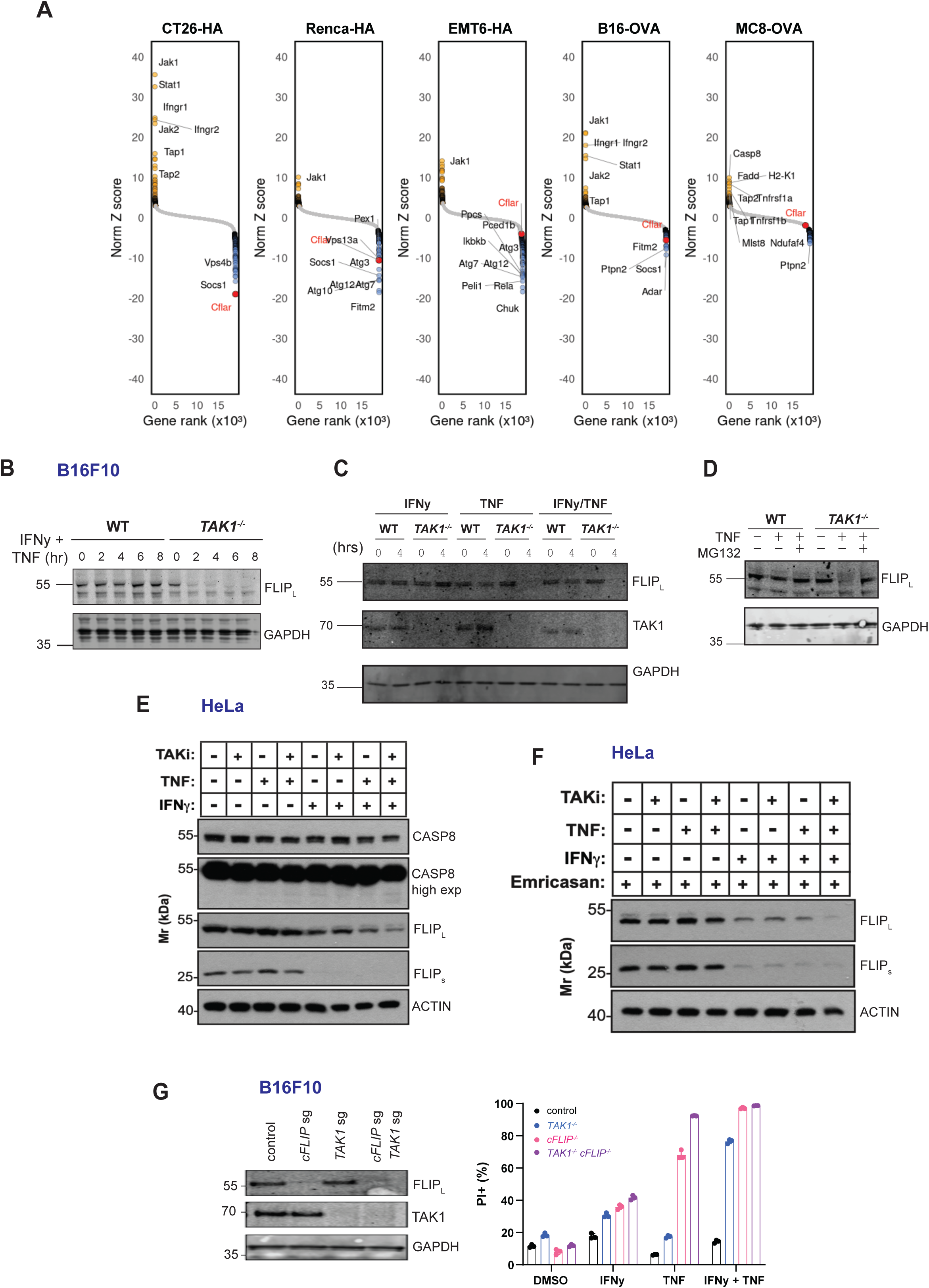
cFLIP inhibits caspase-8 induced cell death in TAK1-deficient cells under inflammatory cytokine exposure. (A) Analysis from Lawson et al (12), highlighting *CFLAR* as an immune evasion gene across multiple cell lines. (B) Time-course immunoblot showing cFLIP (FLIP_L_ isoform) expression in wild-type and TAK1-deficient B16F10 cells treated with IFNγ and TNF for the indicated durations (0-8 hours). (C) Western blot of FLIP_L_ expression after 4-hours stimulation with IFNγ, TNF, or their combination in WT and TAK1-deficient B16F10 cells. (D) cFLIP protein levels in WT and TAK1-deficient B16F10 cells treated with TNF (4 hours) with or without proteasome inhibitor MG132 (5 µM). (E-F) Similar cytokine-induced downregulation of cFLIP (FLIP_L_ and FLIP_s_) observed in HeLa cells (E) treated with IFNy (100 ng/mL), TNF (2 ng/mL) or/and TAKi (5 µM), and where indicated (F), in cells treated with Emricasan (5 µM). All these indicate a conserved effect of TAK1 inhibition across cell types. (G) Flow cytometry quantification of PI⁺ (dead) cells following 24-hours treatment with IFNγ and TNF in control, TAK1-deficient, cFLIP-deficient, or double-deficient B16F10 cells.

To determine whether cFLIP loss was mediated by the proteasome, we treated TAK1-deficient cells with MG132. Indeed, MG132 exposure prevented cFLIP degradation in the presence of TNF (Figure 6D), indicating that cFLIP is actively degraded through a proteasome-dependent mechanism. Similar patterns of cFLIP downregulation were observed in HeLa cells following TNF treatment in the context of TAK1 loss or inhibition (Figure 6E–F). To test the functional relevance of cFLIP degradation, we generated cFLIP knockout cells with or without TAK1 knockout. cFLIP knockout alone modestly increased sensitivity to TNF cytokine, but not IFNy. However, double knockout of TAK1 and cFLIP synergistically increased the cell death (Figure 6G). Collectively, these data identify cFLIP degradation as a key downstream consequence of TAK1 inactivation, linking inflammatory signaling to caspase-8 activation and apoptotic cell death in tumor cells exposed to IFNγ and TNF.

### TAK1 loss sensitizes tumors to T cell–mediated elimination, adoptive cell therapy and correlates with improved immunotherapy response

To evaluate the impact of TAK1 loss on tumor progression in vivo, we implanted control or TAK1-deficient B16F10 cells subcutaneously into immunocompetent C57BL/6 mice and monitored tumor growth and survival (Figure 7A). Western blot confirmed efficient depletion of TAK1 in edited cells (Figure 7B). While TAK1 loss did not affect *in vitro* proliferation (Figure 7C), TAK1-deficient tumors grew significantly more slowly *in vivo* and extended host survival compared to controls (Figure 7D–E), suggesting enhanced immune-mediated rejection. To test whether this tumor suppression was dependent on immune cells, we repeated the experiment in NSG mice, which lack functional lymphocytes and NK cells. In this context, TAK1-deficient tumors grew at rates similar to controls (Figure 7F–G), indicating that the anti-tumor effect of TAK1 loss is immune-dependent.

**Figure 7.**
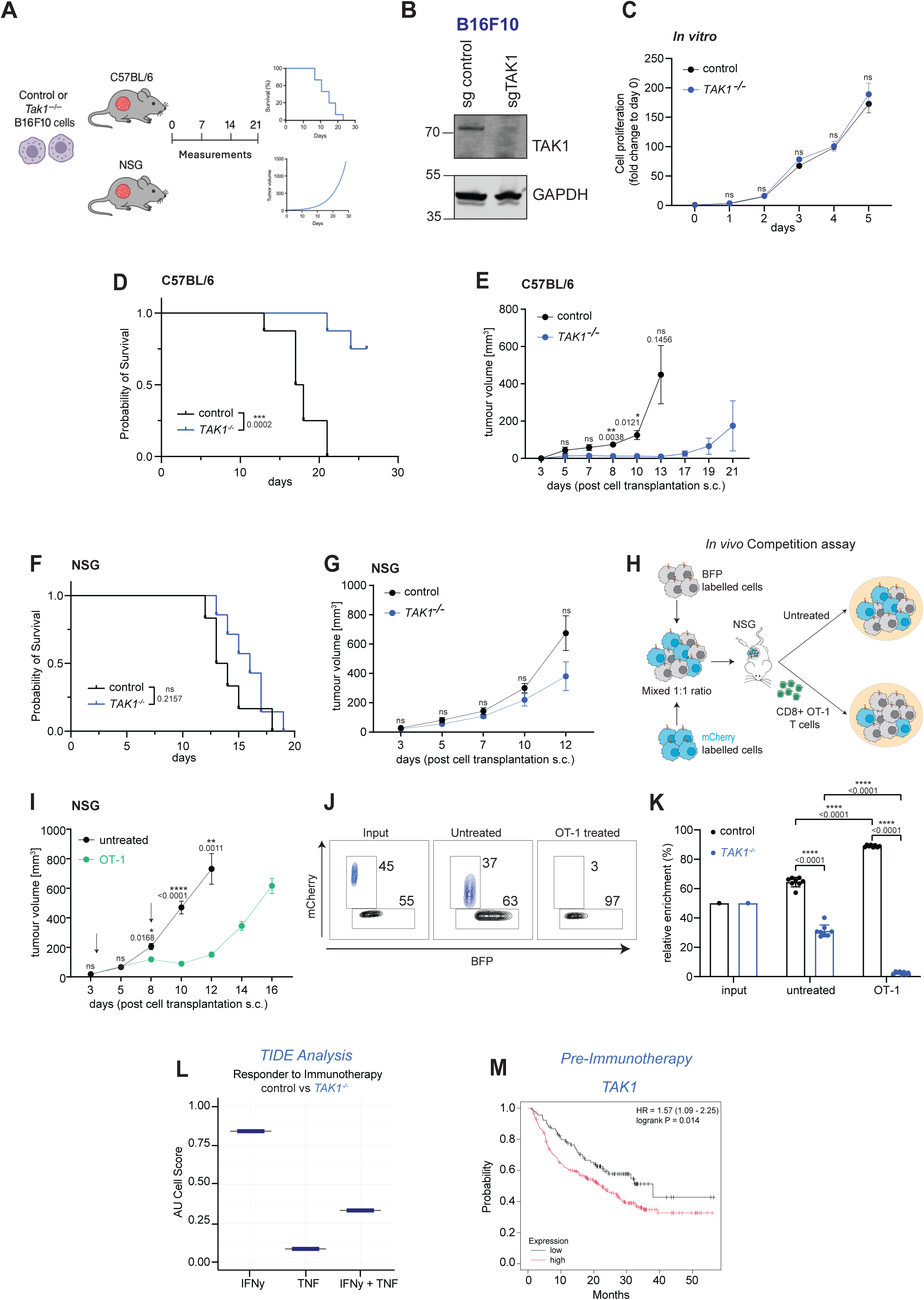
TAK1 deficiency enhances tumor susceptibility to T cell-mediated immunity and is associated with improved cancer immunotherapy response. (A) Schematic of tumor implantation and monitoring protocol. Wild type control or TAK1-deficient B16F10 cells were injected subcutaneously into immunocompetent (C57BL/6) or immunodeficient (NSG) mice, followed by tumor growth and survival analysis. (B) Western blot confirming TAK1 depletion in B16F10 cells transfected with control or Tak1-targeting sgRNAs. (C) *In vitro* proliferation of wild type control and TAK1-deficient B16F10 cells over 5 days. (D–E) Survival curves (D) and tumor growth (E) of WT and *TAK1^-/-^* B16 cells in C57BL/6 mice. Tumour growth data were analysed using ANOVA with multiple comparisons. (F–G) Survival curves (F) and tumor growth (G) WT or *TAK1^-/-^* B16 cells in NSG mice. Tumour growth data were analysed using ANOVA with multiple comparisons. (H) Schematic of *in vivo* competition assay. BFP-labeled WT and mCherry-labeled TAK1-deficient cells were mixed 1:1 and implanted into NSG mice ± adoptive transfer of OT-I CD8⁺ T cells. (I) Tumor volume measurements from competition assay in NSG mice ± OT-I T cells. TAK1-deficient cells were selectively eliminated in the presence of OT-I cells. (J–K) Representative FACS plots (J) and quantification (K) of mCherry⁺ (*TAK1^⁻/⁻^*) and BFP⁺ (WT) tumor cells from *in vivo* competition assays. TAK1-deficient cells are significantly depleted in the OT-I treated group. ****p < 0.0001, ANOVA with multiple comparisons. (L) AUCell scoring using the TIDE algorithm for response to immunotherapy in melanoma using gene signatures derived from the RNA seq data in (Figure 3). (M) Using the algorithm from Kovacs et al (22), we analysed TAK1 expression as a predicative biomarker for response to immunotherapy for melanoma (anti-PD1/antiCTLA-4/combination).

To directly assess T cell-mediated clearance of TAK1-deficient tumors, we conducted an *in vivo* competition assay. WT (BFP-labeled) and TAK1-deficient (mCherry-labeled) B16F10 cells were mixed 1:1 and implanted into NSG mice, followed by adoptive transfer of OT-I CD8⁺ T cells (Figure 7H). In the presence of OT-I cells, tumors were significantly smaller (Figure 7I), and flow cytometry revealed near-complete elimination of TAK1-deficient cells compared to controls (Figure 7J–K). These findings support a model in which TAK1-deficient tumor cells are selectively eliminated by antigen-specific CD8⁺ T cells *in vivo*.

To understand the molecular basis for this enhanced immunogenicity, we performed AUCell scoring derived from the TIDE algorithm (21) for IFNγ and TNF-responsive gene signatures derived from our RNA seq data (Figure 3) in TAK1-deficient cells. TAK1 loss significantly amplified immunotherapy response gene expression upon cytokine exposure, particularly under combined IFNγ and TNF treatment (Figure 7L), consistent with prior in vitro findings of cytokine hypersensitivity. Finally, we examined the clinical relevance of *TAK1* expression in the context of immunotherapy. Using the algorithm from Kovacs et al (22), we analysed *TAK1* expression as a predicative biomarker for response to immunotherapy for melanoma. As can be seen from figure 7M, patients with lower *MAP3K7* expression prior to immunotherapy (anti-PD1/antiCTLA-4/combination) had an improved prognosis. Collectively, these results demonstrate that TAK1 loss sensitizes tumors to T cell–mediated immune pressure and enhances tumor clearance, highlighting TAK1 as a candidate therapeutic vulnerability in immunotherapy-refractory cancers.

## Discussion

While immunotherapies including checkpoint blockade have revolutionized cancer treatment, their efficacy may be limited by tumor-intrinsic survival programs that enable immune escape or prevent cell death. Here, we identify the serine/threonine kinase TAK1 (MAP3K7) as a key tumour intracellular checkpoint that controls whether cells drive inflammation or undergo cell death in response to CTL-derived effector cytokines.

Our findings reveal a pro-survival network in which TAK1 maintains cellular resistance to cytokine-induced apoptosis by maintaining the expression of the caspase-8 inhibitor cFLIP. In the absence of TAK1, and in response to inflammatory stimuli, cFLIP is rapidly degraded via the proteasome, which lowers the threshold for caspase-8 activation and initiates cell death. This process initiates the assembly of a death-inducing complex comprising RIPK1, FADD, and caspase-8. Importantly, this pathway is distinct from canonical perforin-or granzyme-mediated cytolysis (23), which was dispensable for sensitisation to CTL upon TAK1 loss. Instead, TAK1-deficient cells succumb cytokine-dependent form of inflammatory cell death that is initiated by TNF and sensitized by IFNγ. These results uncover a non-canonical, cytokine-driven cytotoxic program that may be broadly relevant in the context of immunotherapy.

Mechanistically, our study integrates and extends prior observations implicating both TNF and IFNγ in tumor clearance (11, 24, 25). While previous work has identified loss of IFNγ signaling as a frequent cause of immune escape and resistance to checkpoint inhibitors (26, 27), our data demonstrate that intact IFNγ signaling can also potentiate tumor cell death when key intracellular checkpoints such as TAK1 are deleted. Similarly, TNF has dual roles in promoting both survival and death (10, 16); we show that the balance tips toward apoptosis in the absence of TAK1, which normally promotes NF-κB– mediated survival downstream of TNFR1 (28, 29). Thus, TAK1 acts as a molecular rheostat that interprets inflammatory cytokine input and dictates tumor cell fate.

Our findings also have direct implications for immunotherapy. TAK1-deficient tumors exhibited delayed growth and enhanced clearance in immunocompetent hosts but not in immunodeficient mice, underscoring the immune-dependent nature of their vulnerability. Moreover, TAK1 loss rendered tumors hypersensitive to adoptively transferred T cells which has implications for chimeric antigen receptor (CAR) T cell therapies. Collectively, our data nominate TAK1 as a novel therapeutic target to augment the efficacy of immunotherapies, especially in tumors with intact antigen presentation but resistance to T cell–derived cytokines.

Our study also raises several key questions. First, the precise post-translational mechanisms by which TAK1 preserves cFLIP stability remain to be defined. Whether TAK1 directly phosphorylates components of the ubiquitin-proteasome system or indirectly regulates cFLIP degradation through NF-κB targets warrants further investigation. Finally, as TAK1 inhibitors are under preclinical and early clinical evaluation in inflammatory diseases, their potential use to sensitize tumors to immunotherapy— particularly when combined with agents that boost T cell cytokine production—represents a promising translational avenue.

In conclusion, we uncover TAK1 as a central node in a tumor-intrinsic survival circuit that shields cancer cells from T cell-derived inflammatory cytokines. By mapping the downstream components of this death pathway—including RIPK1, STAT1, and cFLIP—we provide a mechanistic framework for understanding how inflammatory stress is integrated at the level of tumor cell survival. These insights open new opportunities for therapeutic intervention, particularly in the design of rational combination strategies that disrupt immune resistance and restore effective anti-tumor immunity.

## Acknowledgements

We thank Professor Marco Herold (ONJCRI) for provision of reagents, technology and advice.

## Materials and methods

### Cell cultures and cell line

B16F10 (Murine melanoma cell line; CVCL_0159), CT26 (Murine colon carcinoma cell line; CVCL_7245), MC38 (Murine colon carcinoma line; CVCL_B288), COLO205 (Human colon carcinoma cell line; CVCL_0218), HCT116 (Human colon carcinoma cell line; CVCL_0291) and HEK293T (Human embryonic kidney cell line) cells were cultured in DMEM supplemented with 5% (v/v) Fetal Calf Serum (FCS), 100 U/mL penicillin, 100 mg/mL streptomycin and 2 mM Glutamax at 37°C and 5% CO_2_ incubator. SK-MEL-2 (Human melanoma cell line; CVCL_0062) cells were cultured in RPMI1640 supplemented with 10% (v/v) Fetal Calf Serum (FCS), 100 U/mL penicillin, 100 mg/mL streptomycin and 2 mM Glutamax at 37°C and 5% CO_2_ incubator.

OT-I T cells were obtained by first harvesting spleens from OT-I mice. Following harvest, T cells were stimulated with the chicken ovalbumin peptide_257-264_ SIINFEKL for 5 days before they were used for either cell competition or cell death assays. OT-I T cells were cultured in RPMI1640 supplemented with 10% (v/v) Fetal Calf Serum (FCS), 1% (v/v) sodium pyruvate, 1% (v/v) non-essential amino acids, 100 U/mL penicillin, 100 mg/mL streptomycin, 2 mM Glutamax, interleukin-2 (IL2;100 U/mL) and 0.05% (v/v) 2-mercaptoethanol at 37°C and 5% CO2 incubator.

### Electroporation of Cas9 and sgRNA for generating knockout cell lines

SgRNAs were designed by Synthego and were synthesized by Genescript. The following sgRNAs were mmTAK1 sgRNA_1 (5’ AGATAGAAAGTGAGTCTGAG 3’), mmTAK1 sgRNA_2 (5’ ACACTGTAAACACCAGCTCA 3’), mmTAK1 sgRNA_3 (5’ CCCAGCTTTCAGAATCATGT 3’), hsTAK1 sgRNA_1 (5’ TTAACTCAGGTTGTTGGAAG 3’), mmCASP8 sgRNA_1 (5’ CAATAGCATAAAGACAACTC 3’), mmRIPK1 sgRNA_1 (5’ TCCTGGCCACAGGTACAATG 3’), mmSTAT1 sgRNA_1 (5’ GGTCGCAAACGAGACATCAT 3’), mmCFLIP sgRNA_1 (5’ CTTACCTATAATCAGAAACC 3’), mmIFNGR1 sgRNA_1 (5’ ATTAGAACATTCGTCGGTAC 3’) and mmTNFR1 sgRNA_1 (5’ AGCAGAGCCAGGAGCACCTG 3’). Target cells were electroporated with either a Cas9 only control or Cas9/sgRNA ribonucleoprotein (RNP) complexes using the SF Cell Line 4D-Nucleofector X kit (Lonza). After 5 days, gene knockout was confirmed either by PCR-Sanger sequencing of the indels site (IDT) or by western blotting.

### Gene expression with retroviral vector and lentiviral vector

3xFLAG-(DYKDHDGDYKDHDIDYKDDDK)-tagged wildtype TAK1 (TAK1-WT) and 3xFLAG-(DYKDHDGDYKDHDIDYKDDDK)-tagged kinase inactive TAK1 (TAK1-D175A) were synthesized as gene fragments by Integrated DNA Technologies (IDT) before they were subcloned to either pMSCV-IRES-GFP or pMSCV-IRES-mCherry retroviral vector. Their insertions were confirmed by Sanger sequencing (AGRF). Viral particles expressing the corresponding TAK1 genes were produced in packaging cells (HEK293T) and were used to infect B16F10-OVA. B16F10-OVA cells expressing the corresponding TAK1 protein were selected by sorting GFP or mCherry fluorophore.

FuCas9mCherry lentiviral vector was obtained from Marco Herold. B16F10-OVA. Lentiviral particles were used to infect B16F10-OVA cells as indicated above, and cells were selected by sorting mCherry fluorophore.

### Genome-wide and targeted CRISPR/Cas9 screens

A targeted CRISPR/Cas9 sgRNA screen targeting kinases was performed on B16F10-OVA cells. Cas9 expressing B16F10-OVA were transduced with the kinase targeting CRISPR/Cas9 sgRNA library at multiplicity of infection (MOI) of 0.3 to 0.6. Following transduction, cells were selected with puromycin (1µg/mL) for 6 days and were expanded. Some cells were collected after which T0 time point was taken as a reference. The remaining cells were co-cultured with the SIINFEKL-activated OT-I T cells at an effector to target (E:T) ratio of 1:1. These processes were repeated two more times but with an increasing E:T ratio of 2:1 and 3:1 for 2nd & 3rd round respectively. Surviving cells from each round were snap-frozen and their genomic DNA was extracted using DNeasy Blood and Tissue Kit (Qiagen, Cat#69504). Enriched sgRNAs were amplified using Ex Taq DNA Polymerase (Takara, Cat#RR001B) and purified with Ampure XP beads (Beckman Coulter, Cat#A63880). DNA libraries were run on MiSeq (Illumina) and MaGeCK analysis was conducted to count the reads and perform gene enrichment analysis. The data were then visualized using R package ggplots2.

Another targeted CRISPR/Cas9 sgRNA screen targeting kinases was performed on B16F10-OVA cells. This time, cells were treated with IFNγ (1 ng/mL) or/and TNF (20 ng/mL) for 5 days to validate if *TAK1* is depleted in the surviving cells. Untreated cells after puromycin selection were taken as T0 time point. Enriched and depleted sgRNAs from surviving cells were obtained and analysed with the method described above.

A genome-wide CRISPR/Cas9 sgRNA screen was performed on *TAK1^-/-^*B16F10-OVA cells. Cas9 expressing *TAK1^-/-^* B16F10-OVA were transduced with the genome-wide library as described above. This time, cells were treated with IFNγ (1 ng/mL) and TNF (20 ng/mL) for 4 days. These treatments were repeated three times to enrich cells containing sgRNAs that abrogated the ability of IFNγ and TNF to kill *TAK1^-/-^* B16F10-OVA cells. Untreated cells after puromycin selection were taken as T0 time point reference. Enriched sgRNAs from surviving cells were obtained and analysed with the method described above.

### RNA sequencing preparation and analysis

Total RNA was extracted using the RNeasy Mini Kit (Qiagen, Cat#74106) following the manufacturer’s instructions and quality checked using a 4200 TapeStation machine. 100ng of total RNA was used per sample for NGS library preparation (Illumina TruSeq RNA, Cat#FC-122-2002) as per the manufacturer’s instructions. NGS libraries were quality checked using a 4200 Tapestation, pooled and then sequenced using a P2 100-cycle Illumina sequencing kit (Illumina, Cat#20046811) on the NextSeq 2000 sequencing system with 66bp paired-end reads. RNA-seq reads were aligned to the mouse reference genome GRCm39/mm39 using the Subread aligner (Rsubread v.2.14.2) [BIOINF1, BIOINF2]. Gene-level read counts were obtained by using featureCounts [BIOINF3]. Rsubread inbuilt gene annotation for mm39 was used for read summarization. Genes that failed to achieve a CPM (counts per million) expression value of 1 or greater in at least 3 libraries were excluded from analysis. Read counts were converted to log2-CPM, quantile normalized and precision weighted with the voom function of the limma package [BIOINF4, BIOINF5]. A linear model was fitted to each gene, and the empirical Bayes moderated t-statistic was used to assess differences in expression [BIOINF6]. Genes were called differentially expressed if they achieved a false discovery rate of 5% or less.

### *In vitro* competition and T cell killing assays

For the FACS-based competition assay, wildtype or *TAK1* sgRNA-treated B16F10-OVA cells were equally mixed and plated on a 24 well plate. These cells were previously labelled with unique fluorophore such as mCherry or BFP. Cells were then cultured with either OT-I, supernatants from the OT-I/B16F10 co-culture or cytokines including IFNγ (1 ng/mL) or/and TNF (20 ng/mL) for 2 days. Proportion of live cells within each labelled population was assessed based on BFP or mCherry expression using the BD FACS Symphony Cell Analyzer.

For the FACS-based cell viability assay, cells were cultured with OT-I, IFNγ (1 ng/mL) or TNF (1-20 ng/mL) at various concentrations for 1-2 days. Live and dead cells were suspended in PBS supplemented with 3% (v/v) FBS and 2 µg/mL propidium iodide. Cell viability was measured by assessing the proportion of propidium iodide positive (PI+) cells.

For plate-reader based cell viability assay, luciferase expressing B16F10-OVA cells and OT-1 T cells were used as target cells (T) and effector cells (E), respectively. The E:T ratio was continuously decreased from 2:1 to 1:100. Bioluminescence was measured using the EnSight plate reader (PerkinElmer). D-Luciferin (Thermo Fisher Scientific, Cat# 88291) was added at a final concentration of 70 µg/mL. The percentage of specific lysis was calculated based on the luminescent signal using the following equation: 100*(spontaneous death luminescence – sample luminescence) / (spontaneous death luminescence – maximal killing luminescence).

### Immunoblotting

Whole cell protein extracts were obtained by lysing cell pellets in 1x NuPage LDS (Thermofisher) supplemented with 5% β-mercaptoethanol (β-ME). Cell lysates were boiled for 10 min before proteins were separated on 4-12% Bis-Tris gradient gels (Invitrogen). Afterwards, proteins were transferred onto either PVDF or nitrocellulose membrane using the iBlot2TM dry blotting system (20V, 7 min). The membranes were blocked with 5% skim milk in PBS-Tween (0.1% (v/v)) for 1 hr. Membranes were then probed overnight at 4°C with the following primary antibodies: CASP3 (CST, Cat#9662), cleaved CASP3 (CST, Cat#9661T), CASP8 (CST, Cat#4927T), cleaved CASP8 (CST, Cat#9429T), CASP8, (AdipoGen, Cat#AG-20B-0057-C100), RIPK1 (CST, Cat#3493S), PARP (CST, Cat#9542T), STAT1 (CST, Cat#9172T), TAK1 (CST, Cat#5206S), CFLIP (CST, Cat#56343), A20 (Santa-Cruz, Cat#sc-166692), FADD (CST, Cat#2782), TRADD (Santa-Cruz, Cat#sc-46653), ACTIN (Proteintech, Cat#66009-1-Ig) and GAPDH (Invitrogen, Cat#437000). The next day, membranes were washed with PBS-T in between incubation before addition of IR800-or Alexa Flour 680-conjugated secondary antibodies (Li-Cor) for 1 hr at 25 °C. Proteins were detected by scanning membrane on Odyssey Imaging System (Li-Cor) and was visualized using Affinity Photo and Adobe Illustrator.

### CASP8 Immunoprecipitation

HeLa cells were treated with IFNy (100 ng/mL), TNF (5 ng/mL) or/and Takinib (5 µM) in the presence of Emricasan (5 µM) for 24 hours. After treatment, cells were first washed with 5mL of ice-cold PBS and lysed with 1 mL IP lysis buffer including 150mM NaCl, 2mM EDTA, 1% Triton-X, 10% glycerol, 0.1 mM PMSF, 0.4mg/mL Aprotinin and 10mg/mL leupeptin,1mM iodoacetamide and 5mM PR-619. Then, cells were scraped and collected into tube and pelleted at 15000g for 20 min at 4°C followed by transfer of supernatants into new tubes. Then, lysates were pre-cleared using Protein A/G PLUS-Agarose beads (sc-2003) for 1h at 4°C. Then, 50 mL of input samples were taken from each group, followed by immunoprecipitation of CASP8 by adding fresh Protein A/G PLUS-Agarose beads and 1mg of Anti-CASP8 IP antibody (MBL, Cat#M058-3) per tube for 4h at 4°C. At the end of the incubation, beads were centrifuged at 1000g for 1 min and washed three times using 1mL of IP lysis buffer. Lastly, samples were eluted from the beads with 100mL of SDS lysis buffer. Cell lysates and IP samples were then subjected to immunoblotting.

### Cytotoxic lymphocyte killing assays

Luciferase-expressing tumor cells were generated by lentiviral transduction using an FUL2tG expression construct driving co-expression of luciferase and mCherry. Virus was generated by overnight transfection of HEK293T cells using calcium phosphate precipitation2 with plasmid DNA alongside packaging plasmids (pMDL, and pRSV-Rev) and VSV-G envelope plasmid (pMD2.G). After 48 hours, virus-containing supernatant was collected, filtered (0.45 μm), and supplemented with polybrene (8 μg/mL) before adding to MC38-OVA and B16-OVA tumor cells overnight. One week after transduction, cells were sorted for mCherry expression via fluorescence-activated cell sorting (FACS) using a FACSAria Fusion flow cytometer (BD Biosciences). Cell killing was assayed through addition of luciferin and emitted luminescence measured with a luminometer.

### Phosphoproteomics

Enrichment: Enrichment of phosphorylated peptides was based off the EasyPhos Protocol (30). In brief, cell pellets were lysed in chilled SDC lysis buffer (4% (w/v) sodium deoxycholate and 100 mM Tris-HCl, pH 8.5) and immediately heat-treated at 95°C for 5 minutes. Cells were sonicated with a tip probe, clarified, and protein amounts were adjusted to 1 mg using a bicinchoninic acid (BCA) assay (1:50 ratio of CuSO4:bicinchoninic acid). Proteins were then reduced with 10 mM tris(2-carboxy(ethyl)phosphine) (TCEP) and alkylated with 40 mM 2-chloroacetamide (CAA) for 5 minutes at 45°C with shaking (1500 rpm). Proteins were digested overnight to peptides with trypsin and lysC resuspended in 0.1% acetic acid (1:160, enzyme:protein, w/w) at 37°C with shaking (1500 rpm). Isopropanol (final concentration 50%), trifluoroacetic acid (TFA, CF3CO2H, final concentration 6%) and monopotassium phosphate (KH2PO4, final concentration 1 mM) was added to the peptides and vortexed. Peptides were clarified at 16,000 g for 5 minutes and supernatant was transferred to a deep-well plate containing 12 mg of TiO2 beads in 100% acetonitrile (ACN) and shaken at 45°C and 1,500 rpm for 5 minutes. Samples were spun at 2,000 g for 1 minute to settle beads, and supernatant was aspirated. Beads were washed five times with 5% (v/v) TFA and 60% (v/v) isopropanol and transferred to C8 Stage Tips (prepared in house) in 0.1% (v/v) TFA and 60% (v/v) isopropanol and spun at 1,500 g to dryness. Phosphopeptides were then eluted off beads and through stage tip with 32% (v/v) acetonitrile and 5.6% (w/v) ammonium hydroxide (NH4OH) and centrifugation at 1,500 g. Samples were dried on a SpeedVac at 45°C and resuspended in 0.1% (v/v) TFA and 99% (v/v) isopropanol and transferred to the top of a SDB-RPS Stage Tip (prepared in house). Stage Tips were spun to dryness (1,500 g) and washed with 0.1% (v/v) TFA and 99% (v/v/) isopropanol followed by a wash with 0.2% (v/v) TFA and 5% (v/v) ACN. Peptides were eluted with 0.14% (w/v) ammonium hydroxide in 60% (v/v) ACN, dried to completeness at 45°C on the SpeedVac, and resolubilised in mass spectrometry loading buffer (2% (v/v) ACN and 0.1% (v/v) formic acid). Mass spectrometry: The peptides were separated using reverse-phase liquid chromatography on a 15 cm C18 fused silica column with an integrated emitter tip (IonOpticks, ID 75 µm, OD 360 µm, 1.6 µm C18 beads) on a Thermo Scientific NeoVanquish LC coupled to Thermo Scientific Orbitrap Astral. Peptides were analysed on a 23 min linear analytical gradient of increasing buffer B (80% ACN, 0.1% FA) from 2 to 34%. Data was acquired in a data independent (DIA) mode. The MS1 settings were as follows: Orbitrap resolution 240,000; scan range m/z 380-980; AGC target 500%. The DIA parameters were as follows: isolation window: 4; HCD collision energy: 25%; scan range: m/z 145-1450; maximum injection time: 6 ms; AGC target: 800%. Raw DIA data were analysed on Spectronaut (31) version 19.9 using BGS factory settings of directDIA analysis with serine/threonine/tyrosine phosphorylation added as a variable modification (which included cysteine carbamidomethylation as fixed modification and N-terminal acetylation and methionine oxidations as variable modifications). PTM localization filter was selected and minimum localization threshold was set to zero. Result filters m/z was set to 1800 (max) and 300 (min) with a relative intensity set to 5. DIA analysis identification had all precursor Qvalue cutoff, precursor PEP cutoff, protein Qvalue cutoff (experiment and run), and protein PEP cutoff set to 0.01. Mouse (Mus musculus) reference proteome was used for database searching. All data analysis of the phosphoproteomics was performed using Perseus software (version 1.6.2.3) (32) and subsequently visualized in R. For phosphosite analysis, Spectronaut normal report output tables were collapsed to phosphosites and the localization cutoff was set to 0.75 using the peptide collapse plug-in tool [4] that collapses phosphoions to phosphosites. Intensities were log2-transformed and filtered for at least 100% valid values in one condition of 4 biological replicates. Missing values were imputed based on a normal distribution (width 0.3. down shift 1.8). A student’s T-test (two-sided) (FDR = 0.05) was performed between TNF and IFNy treated and untreated conditions.

### *In vivo* tumour models

B16F10 cells were electroporated with control sgRNA or sgRNA targeting *MAP3K7*. 5 days later, cells were analyzed by western blot for TAK1 expression. Furthermore, 2 x 10^^5^ cells were injected subcutaneously into recipient C57/BL6 mice or NSG mice (n = 10/group). Tumor growth was measured every 2-3 days using a callipers. Mice were euthanised when tumors reached an ethical endpoint of 500mm^3^. All *in vivo* experiments performed in this study were performed using procedures approved by the Austin Ethics Committee.

---

**Supplementary Figure 1.**
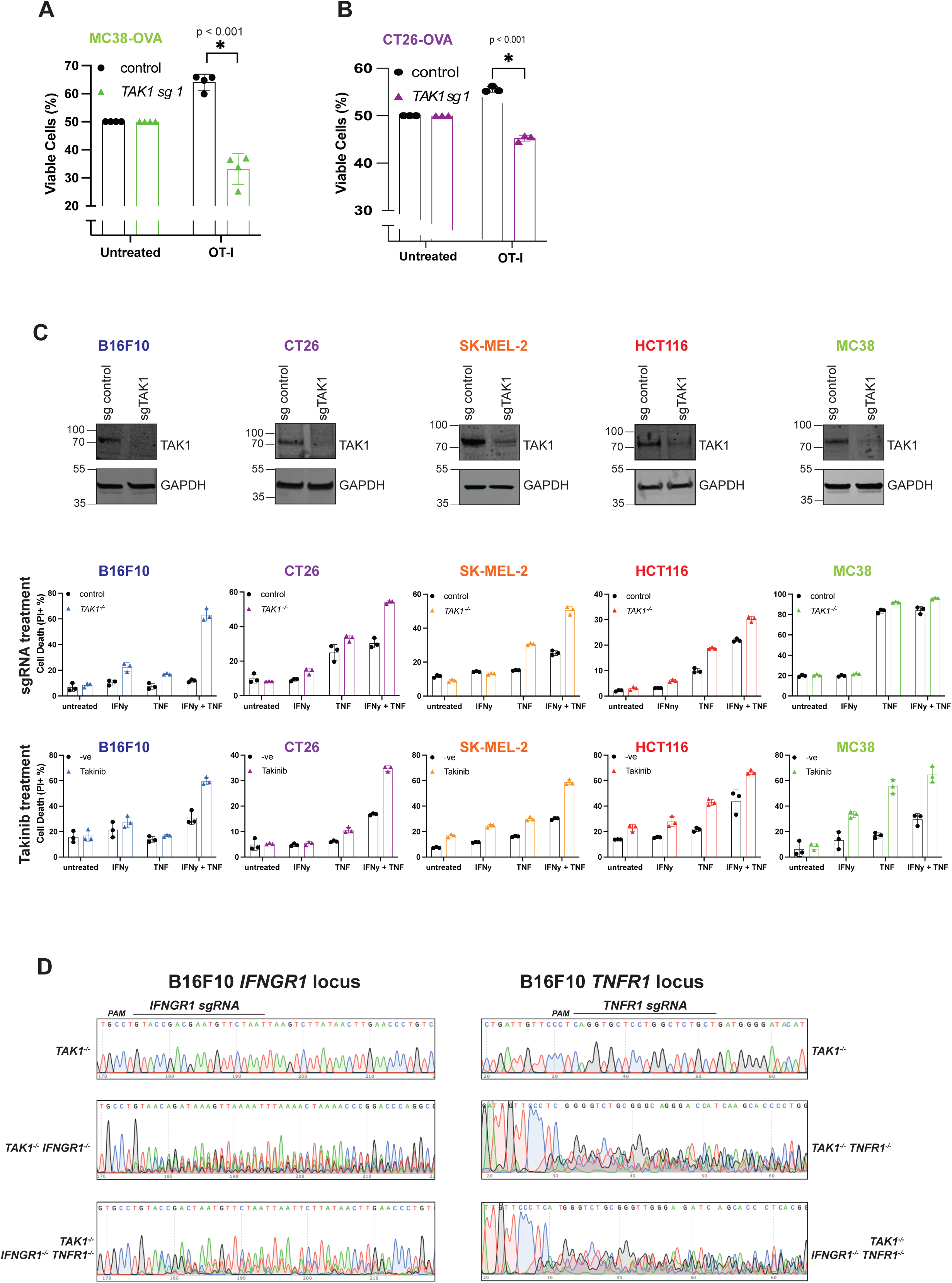
Inactivation of TAK1 is required for many cells to undergo IFNy and TNF induced killing. (A-B) Wild type or TAK1-deficient MC38-OVA (A) / CT26-OVA (B) were labelled with mCherry or BFP and subject to a competition assay under OT-I CD8+ T selection pressure (E:T ratio 1:1) as described in Figure 2A. Relative proportions of WT/KO cells were analysed by FACS after 48 hours. (C) The indicated cell lines (B16F10, CT26, SK-MEL-2, HCT116, MC38; WT or *TAK^-/-^*) were treated with IFNy (1 ng/mL), TNF (1-20 ng/mL) or both for 24-48 hours followed by PI uptake measurement using flow cytometry (middle panel). The same cytokine treatment assay was performed on wild type cells in the presence or absence of Takinib (10 µM) (bottom panel). (D) Sanger sequencing trails of the indicated sgRNA target sites for the indicated genes (*IFNGR1*, *TNFR1*) to confirm CRISPR-mediated deletion of cells used in Figure 2N.

